# First detection of herpesvirus and prevalence of mycoplasma infection in free-ranging Hermann’s tortoises (*Testudo hermanni*), and in potential pet vectors

**DOI:** 10.1101/2021.01.22.427726

**Authors:** Jean-Marie Ballouard, Xavier Bonnet, Julie Jourdan, Albert Martinez-Silvestre, Stéphane Gagno, Brieuc Fertard, Sébastien Caron

## Abstract

Two types of pathogens cause highly contagious upper respiratory tract diseases (URTD) in Chelonians: testudinid herpesviruses (TeHV) and a mycoplasma (*Mycoplasma agassizii*). In captivity, these infections are frequent and can provoke outbreaks. Pet trade generates international flow of tortoises, often without sanitary checking; individuals intentionally or accidentally released in the wild may spread pathogens. A better understanding of the transmission of infectious agents from captivity to wild tortoises is needed. Many exotic individuals have been introduced in populations of the endangered western Hermann’s tortoise (*Testudo hermanni hermanni*), notably spur-thighed tortoises (*Testudo graeca*). We assessed the presence of TeHV and mycoplasma in native western Hermann’s tortoises and in potential pet vectors in south-eastern France. Using a large sample (N=572 tortoises), this study revealed, by PCR, the worrying presence of herpesvirus in 7 free-ranging individuals (3 sub-populations). Additionally, *Mycoplasma agassizii* was detected, by PCR, in 15 of the 18 populations sampled with a frequency ranging from 3.4% (1 of 29 tortoises) to 25% (3 of 12 tortoises). Exotic spur-thighed tortoises showed high frequency of mycoplasma infection in captivity (18.2%) and in individuals (50%) found in native Hermann’s tortoise sub-populations, suggesting that this species could be a significant vector. The paucity of information of TeHV on European tortoise’ URTD in natural settings, especially in combination with mycoplasma, prompts for further studies. Indeed, sick tortoises remain concealed and may not be easily detected in the field. Our results indicate a good health for most infected tortoise but it should be screened in the field as well as in captivity.

## Introduction

Emerging infectious diseases (EIDs) represent a growing challenge for biodiversity conservation (Daszak et al., 2000; Deem et al., 2001). During the last decades, rapidly spreading diseases are suspected to have wreaked havoc worldwide among amphibians and reptiles (Daszak et al., 2000; Tompkins, et al., 2015). In tortoises, *Testidinid herpesviruses* (TeHV) and *Mycoplasma spp*. are two dangerous pathogens for wild populations (Origgi, 2012; Marenzoni et al., 2018). Both are highly contagious and are involved in the Upper Respiratory Tract Disease Syndromes (URTD); infection with these pathogens can provoke illness entailing high morbidity and mortality (Brown et al., 1994; Goessling et al., 2019). For example, in the 80’s, mycoplasma epizooties were responsible of multiple collapses of desert tortoise populations in North America (Jacobson et al., 1991; Brown et al., 1994). Currently, several tortoise species such as Gopher tortoise (*Gopherus agassizii*), or captive spur-thighed tortoise (*Testudo graeca*) are impacted by URTD (Marschang and Schneider, 2007; Weitzman et al. 2017). Monitoring the health status of free-ranging chelonians, with a focus on two major agents causing URTD, is thus a conservation priority.

On the other hand, global trade of reptiles is flourishing; tens of thousands of individuals from an increasing number of species are displaced among continents under minimal (or non-existent for illegal trade) sanitary monitoring (Auliya et al., 2016). The resulting flows of individuals open major routes for the expansion of EIDs (DiGeronimo et al., 2019). Pet tortoises host both TeHV and *Mycoplasma spp*. (Martínez-Silvestre et al., 1999; Sandmeier et al., 2009; Lecis et al., 2011; Salinas et al., 2011; Origgi, 2012). The primary route of transmission of herpesvirus and mycoplasma is believed to be horizontal via contact between individuals (DiGeronimo et al., 2019). Interspecific transmission has been demonstrated (Origgi et al., 2004; Soares et al., 2004; Salinas et al., 2011). Tortoise species are often mixed up in cages or enclosures. Overall, very high prevalence of infection in captivity is expected (Kolesnik et al., 2017). Unfortunately, captive individuals are frequently released into the wild, intentionally or accidentally, where they can settle in novel habitats while carrying pathogens that may threaten native populations of tortoises (Sandmeier et al., 2009; Jacobson et al. 2014; Whitfield et al., 2018). This process opens highways for pathogens. The scarcity of investigations in natural populations, except in USA, means that possible role of pet tortoises as agents of contamination is not quantified in Europe, as in most parts of the world (Jacobson and Berry, 2012; Kane et al., 2017; Orton et al., 2020).

Fragile inbred populations are particularly at risk. Reduced phenotypic diversity and genetic depression often hinder physiological and demographic resistances to diseases (Frankham et al. 2002; Spielman et al., 2004). Many species are subjected to all the threats above, including the not yet evaluated risk of simultaneous contamination by TeHV and mycoplasma. The Hermann’s tortoise provides a typical example of such a situation. Two Hermann tortoise subspecies are currently recognized (Pérez et al., 2013): the Western Hermann’s tortoise (WHT, *Testudo hermanni hermanni*) that occurs west of the Po Valley in Italy (e.g. Italian Peninsula, Sardinia, Corsica, southeastern France, northeastern Spain) and the Eastern Hermann’s tortoise T. h. boettgeri (EHT) found in Mediterranean regions of the Balkan Peninsula and in small islands spread in the eastern Mediterranean sea. The two subspecies come into contact in north eastern Italy where they possibly hybridize naturally (Pérez et al., 2013). The WTH is severely threatened by habitat loss and fragmentation, frequent fires, illegal harvesting and predation by feral animals. As a result, continental populations drastically decreased (Livoreil, 2009). Previously abundant in continental southeastern France, relict isolated WHT sub-populations persist in the Massif des Maures (Var district, 83) and in adjacent plains (Livoreil, 2009; Bertolero et al., 2011). Delayed maturity, low fecundity and low population turn over mean that it is particularly sensitive to a decrease of adult survival (Bertolero et al., 2011).

Intensive and long-lasting legal and illegal pet trades (CITES 2014) provoked substantial introgression of EHT and of other tortoise species (e.g. spur-thighed tortoise) inside the natural repartition area of WTH (Martínez-Silvestre et al., 2001), while various species originating from other continents are sporadically found in the wild (unpublished data). These exotic tortoises easily breed in captivity and are frequently owned as pets; they occur in many properties spread across the entire (remaining) distribution range of the native WHT. A considerable pool of captive individuals from various uncontrolled origins strongly enhances the likelihood of contacts with free-ranging native WHT.

Cross-species transmissions have been documented both for TeHV and mycoplasma; experiments demonstrated that these pathogens can infect Hermann and spur-thighed tortoises (Origgi et al., 2004; Soares et al., 2004; Salinas et al., 2011). In Europe, TeHV-1 and −3 affect most of the Testudinidae species raised in captivity (Origgi, 2012; Marschang and Schneider, 2007). More generally, approximately 48% of individuals belonging to various terrestrial and aquatic pet chelonians were positive for herpesvirus or mycoplasma while a positive correlation was observed between the two pathogen detection frequencies (Kolesnik et al., 2017). Like TeHV, mycoplasma was detected in tortoises kept in captivity and in outdoor enclosures in Europe (Lecis et al., 2011; Salinas et al., 2011).

Mortality due to TeHV and mycoplasma is well documented in wild tortoises in USA (Jacobson et al., 2012, 2014), but only in captivity in European tortoises (Soares, 2004; Kolesnik, et al., 2017). In the latter, mortality rate caused by TeHV is higher in Hermann’s tortoise compared to other species, suggesting a recent and more deleterious contact between the host and the pathogen (Soares et al., 2004). Possible occurrence and consequences of co-infection by TeHV and mycoplasma are not documented, at least in the Hermann’s tortoise. Both frequency and possible severity of TeHV and mycoplasma infections remain unexplored in native populations of tortoises in Europe. Overall, possible impact of worrying EIDs has not been assessed in natural setting in the Mediterranean basin that hosts many endemic tortoises (Gracià et al., 2020). This issue is urgent because exogenous tortoises are frequently observed in the wild where they may carry new pathogens (Lecis et al., 2011; Hidalgo-Vila et al., 2020).

Accurate monitoring of the frequency of infections with these pathogens and information about the prevalence of URTD in the remaining populations of WHT is thus needed, notably because these infectious agents can induce deleterious chronic diseases that are not easily diagnosed in long-lived organisms (Sandmeier et al., 2013). This study reports results from the first comprehensive survey across the distribution range of the WHT in continental France.

## Methods

### Tortoise sampling

From 2012 to 2016, 18 sites were monitored covering most of the distribution area of the WTH subspecies in continental France (besides these surveys, individuals were also opportunistically sampled throughout the distribution area) (Figure 1; Livoreil, 2009). Free-ranging tortoises are cryptic; thus, in addition to visual searching, trained dogs were used (Ballouard et al., 2019). Surveys were performed during the activity season of the species (from March to October), mostly in spring (111 searching days in spring, 34 in summer and 15 in autumn). All tortoises sighted were captured: 457 free-ranging individuals (421 WHT and 36 exotic specimens) were sampled (25.5 tortoises per site on average). In addition, 95 captive (pet) tortoises were sampled in 21 different properties in surrounding areas (5.2 individuals per property on average). Finally, 20 vagrant isolated individuals found in urban, or peri-urban areas, were also tested; likely they were pets intentionally released or that escaped from gardens. Most tortoises were adult (96%) and sex ratio was balanced (281 females, 267 males, 24 immatures). Overall, with respect to sampling context, we obtained three categories of individuals: a) free-ranging, b) captive (pet), and c) vagrant (N total=572). Each sampling category contained both native and exotic tortoises.

**Figure 1.**
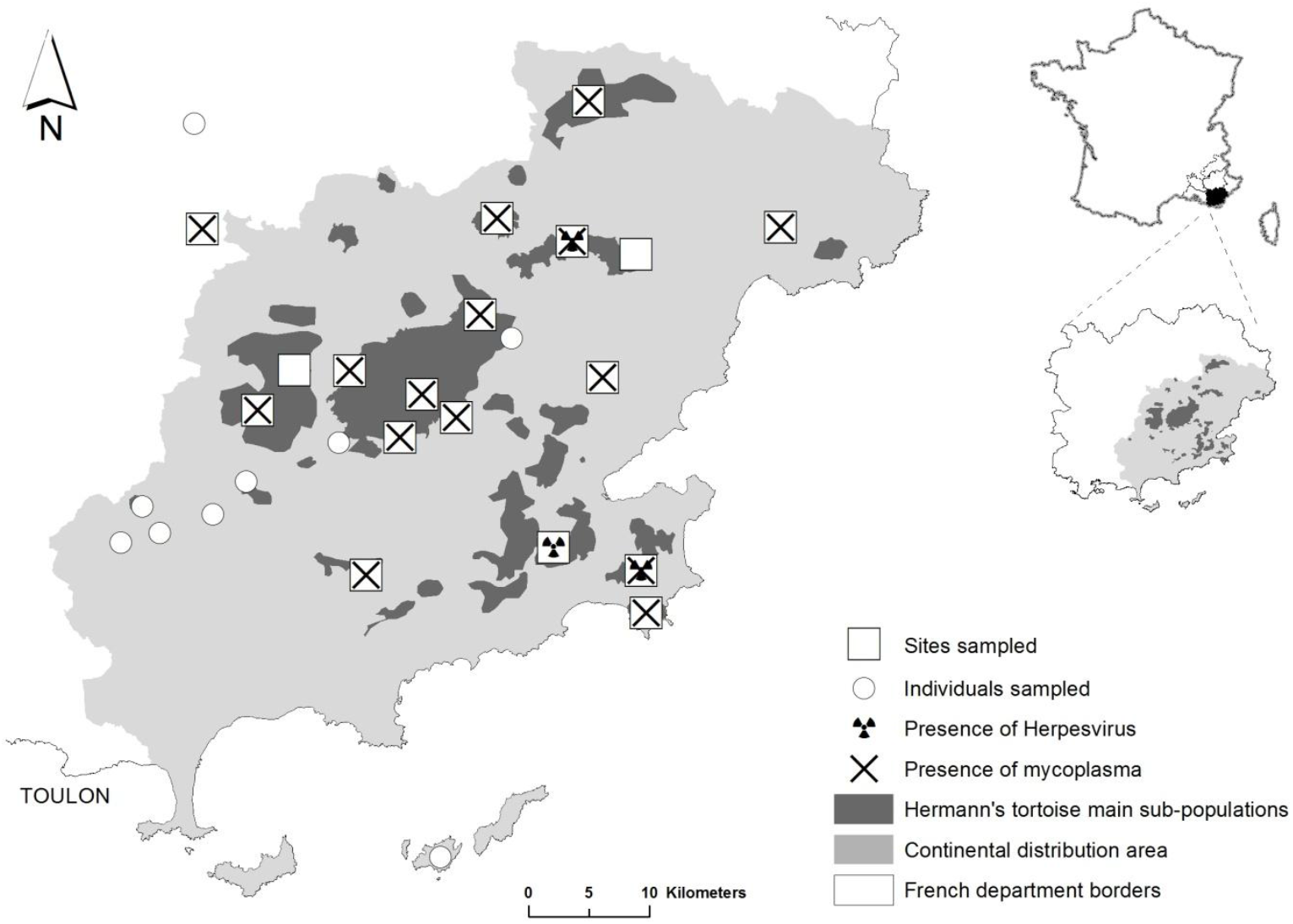
Sampling sites inside the French continental distribution range of WHT.

Individuals were assigned to species (e.g. *T. graeca vs. T. hermanni*) or subspecies (WHT vs. EHT) according to their morphological characteristics. Testudo species are easily distinguished (Bertolero et al. 2011), subspecies not. The following criteria were used to discriminate WHT from EHT: yellow subocular scales, black continuous plastral bands, narrow vertebral scute, supracaudal scute divided, long corneous tip of the tail, corneous tubercles on the inner side of the tight, ratio of pectoral vs. femoral seams (Bertolero et al., 2011; Soler et al., 2012). Hybrids WHT x EHT displayed various combinations of phenotypic characters and could not be identified with certainty (especially F2, unpublished genetic results). Easily identifiable hybrids were brought to the Soptom rescue center.

Each tortoise was measured by strait carapace length (SCL), sexed when possible (small immatures cannot be easily sexed), and weighted to the nearest gram. Individuals larger than 100 mm in SCL were considered adult. Following blood, oral and nasal epithelium sampling, individuals were subjected to clinical inspection (see below), and then they were released at the place of capture, generally within 30 min. To ensure that researchers did not spread pathogens and did not contaminate samples, they cleaned their hands and clothes using Vircon spray 1% (Bayer©); equipment was cleaned with alcohol. Samples were stored using one box per site.

Most individuals examined were free-ranging WHT (N=421; Table1). We identified 11 EHT: 2 free-ranging individuals introduced into WHT populations, 8 captive (pets), and one vagrant. Thirty nine WHT x EHT easily identifiable hybrids were observed: 24 free-ranging, 9 captive and 6 vagrants. Twenty four spur-thighed tortoises were examined: 10 of them were free-ranging and thus have been introduced into WTH populations, 12 were found in private properties (pets), and 2 were vagrant. Finally, 1 captive marginated tortoise (*Testudo marginata*) was sampled.

**Table 1.** Numbers of individual tortoises sampled in function of species and subspecies (taxon), and in function of the situation where the animals were found (free-ranging, captive or vagrant, see text for details). *Testudo hermanni hermanni* is coded WHT, *T.h. boettgeri* EHT. WHT x EHT stands for hybrids. Different subspecies of the spur-thighed tortoises (T. graeca) are not distinguished.

|  | Free-ranging | Captive (pet) | Vagrant | TOTAL |
| --- | --- | --- | --- | --- |
| *T. h. hermanni* | 421 | 65 | 11 | 497 |
| *T. h. boettgeri* | 2 | 8 | 1 | 11 |
| *T. g. ibera* | 1 | - | - | 1 |
| *T. g. sp* | 9 | 12 | 2 | 23 |
| *T. marginata* | - | 1 | - | 1 |
| WHT x EHT | 24 | 9 | 6 | 39 |
| TOTAL | 457 | 95 | 20 | 572 |

### Tissue sampling

Blood (0.4 - 0.7 ml) was collected from the subcarapacial plexus (Hernandez-Divers et al. 2002) using 1 ml syringes (Injekt-F – B Braun) and sterile needles (26G to 27G, Terumo Neolus, adjusted to the size of the animal). Subcarapacial plexus delivers various mixtures of blood and lymph, especially using needles larger than 27G (Bonnet et al. 2016); we did not notice such mixture during sampling. Blood was immediately placed in Sodium or Lithium heparin. Samples were gently homogenized and stored (max 4 hours) in ice-cooled containers until centrifugation (1500 rpm for 5 min). Plasma was stored at −25°C until analyses. Aliquots were distributed in two tubes: 50 μl for microbiological analysis and the rest for biochemical analyses.

We sampled oral and nasal epitheliums and mucus. We injected ~ 0.5 ml of sterile saline (0.9% sodium chloride, Lavoisier) to flush the nasal cavity. The resulting fluid was collected with a syringe (0.1 ml) in each nostril and immediately stored in a 0.5 ml sterile conical tube. Oral samples were collected with a brush (Cervibrush + LBC, Endocervical sampler, CellPath) inserted inside the oral cavity: choana and mucosal surfaces of the tongue and of the beak were targeted. Brushes were stored individually in a 0.5 ml sterile conical tube containing 0.3 ml of sodium chloride (0.9%) to avoid desiccation of the mucus. All samples were placed in ice-cooled containers in the field and stored at −25°C until analyses.

All samples were shipped frozen to Staaliches VetUAmt laboratory in Detmold, Germany, for analyses.

### Pathogen screening

The presence of Testudinid herpesvirus (TeHVtype 1 and 3) was assessed using both polymerase chain reaction (PCR) (see method in Teike et al., 2000) and induced antibody responses by serum neutralization test (SN) (Soares et al., 2004; Salinas et al., 2011; Origgi, 2012). PCR test can be applied to TeHV-1 and TeHV-3 in any species of tortoise (Salinas et al. 2011). A PCR test was considered positive for TeHV when DNA of pathogens was detected in oral mucus. However, the mucus of individuals that do not present clinical signs is generally less rich in viral DNA compared to the mucus of individuals displaying clinical signs (Origgi, 2012). In addition, false negatives have been observed, likely due to sampling methodology difficulties or to oscillating elimination of some viruses such as herpesviruses (Marschang 2019). Thus, we used SN testing as a complementary method to detect the presence of TeHV types 1 and 3 circulating antibodies on plasma aliquots (Origgi, 2001, 2012).

The presence of *Mycoplasma agassizii* was assessed using PCR (see method in Brown et al. 1999), no serological test being available in Europe during the study (2012-2016). Thus, PCR were used to detect active infection by mycoplasma and TeHV (Soares et al., 2004).

### Clinical inspection

All tortoises were visually inspected for clinical symptoms of URTD disease that are usually associated with mycoplasma and herpesvirus infections (Jacobson et al., 2014): keratitis, conjunctivitis, ocular and palpebral oedema, and nasal discharges (mucopurulent oculonasal discharge), necrotic spots on the oral mucosa and on the tongue, rhinitis, lethargy and low body condition (Brown et al., 2002; Berry and Christopher, 2001; Sandmeier et al., 2009).

## Results

Many tortoises were tested for both pathogens. Table 2 provides the numbers of tests performed on the different sampling categories (e.g., free-ranging, captive) and tortoise species. PCR testing revealed that seven free-ranging WHT (six adult females and one adult male) were TeHV positive (2.8 %, n=7/253), and that 3 sub-populations (i.e. sites) were concerned (Tables2 and 3). SN tests for TeHV were all negative (Table2).

**Table 2.**
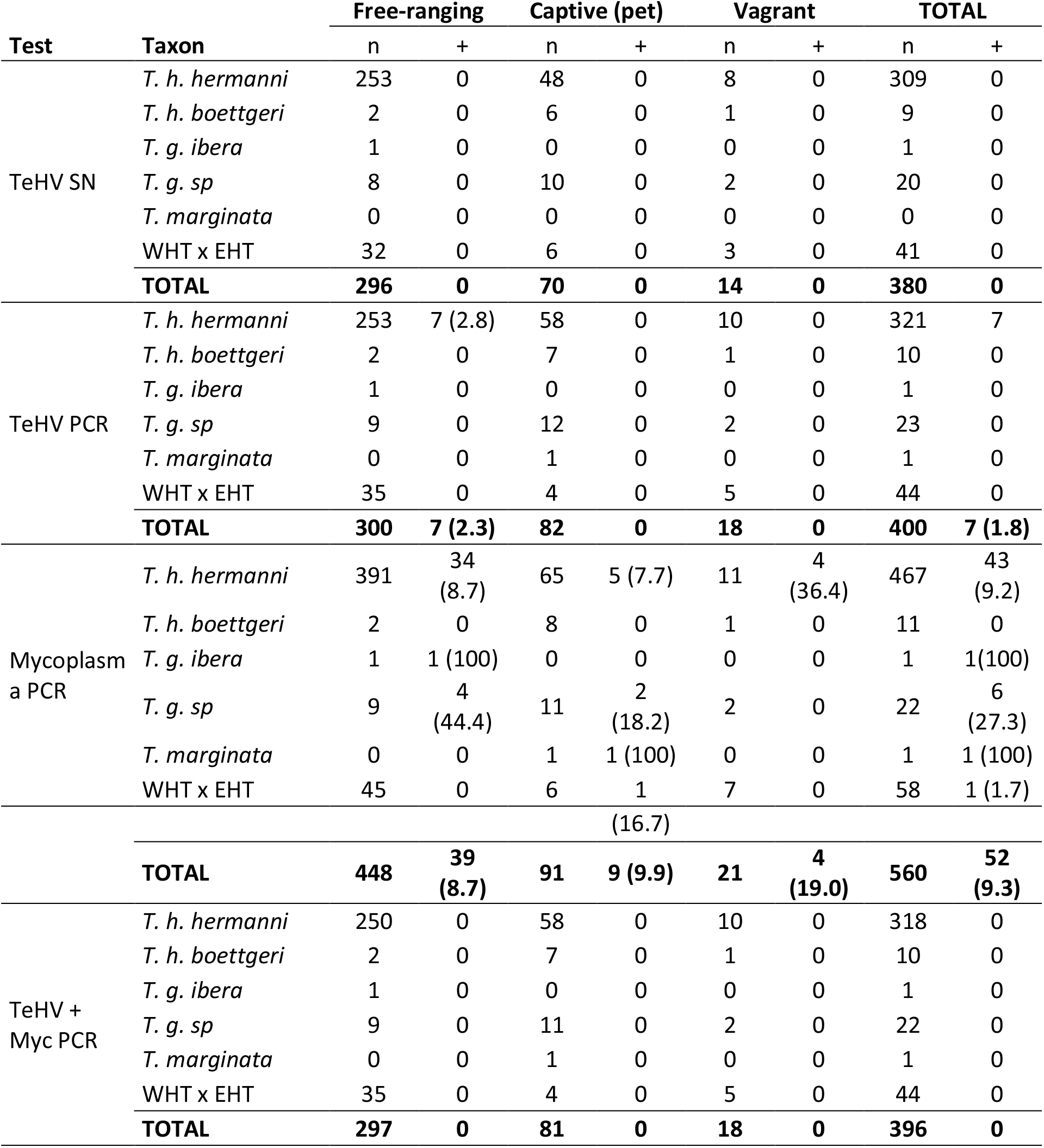
Results from the PCR tests for herpesvirus and mycoplasma infections according to taxon and in function of the situation where the tortoises were found. TeHV stands for Herpesvirus, SN for Serum Neutralization, PCR for polymerase chain reaction test, Mycoplasma for *Mycoplasma agassizii*. Most individuals were tested for both pathogens with PCR (TeHV + Myc). Sample size is provided (n); the sign + indicates the number of positive tests associated. Percentages of positive tests are provided in brackets.

**Table 3.** Results from the PCR tests for herpesvirus (TeHV) and mycoplasma (Mycoplasma) infections of free-ranging tortoises in the 18 sites sampled. Each site hosts a free-ranging population of WHT and often exotic tortoises. Other stands for tortoises opportunistically sampled in various locations. Numbers of individuals sampled and tested positive in each site are provided (n, +). The percentage of positive tests is indicated into brackets.

| Sites | TeHV |  | *Mycoplasma* |  |
| --- | --- | --- | --- | --- |
|  | n | positive results | n | positive results |
| 3 caps | 12 | 0 | 18 | 2 (11.1) |
| La Pardiguière | 28 | 0 | 28 | 1 (3.6) |
| Callas | 17 | 0 | 24 | 4 (16.7) |
| Carcès | 10 | 0 | 10 | 2 (20) |
| Cogolin-La Môle | 20 | 4 (20) | 23 | 0 |
| Estérel | 18 | 0 | 28 | 5 (17.9) |
| Flassans calcaire | 13 | 0 | 21 | 0 |
| Lambert | 15 | 0 | 23 | 1 (4.3) |
| Le Muy - St Luen | 12 | 1 (8.3) | 21 | 2 (9.5) |
| Les Arcs | 13 | 0 | 12 | 3 (25) |
| Neuf riaux / Est PDM | 11 | 0 | 29 | 1 (3.4) |
| Ramatuelle | 9 | 2 (22.2) | 16 | 1 (6.3) |
| Redon | 18 | 0 | 24 | 2 (8.3) |
| Reserve National des Maures | 18 | 0 | 21 | 3 (14.3) |
| Roquebrune sur Argens | 11 | 0 | 17 | 0 |
| Les Mayons | 25 | 0 | 60 | 5 (8.3) |
| Sainte-Maxime | 15 | 0 | 18 | 3 (16.7) |
| Vidauban | 19 | 0 | 35 | 2 (5.7) |
| Other | 16 | 0 | 20 | 2 (10) |
| **TOTAL** | **300** | **7 (2.3)** | **448** | **39 (8.7)** |

A total of 52 individuals (species and subspecies pooled) were positive for mycoplasma DNA (prevalence Pr=9.2%) (Table2). The proportion of individuals infected was not influenced by sampling categories (χ2= 1.56; df=2; p=0.45). We found a significant effect of the species on the observed prevalence of mycoplasma infection (χ2= 20.41; df=3; p<0.001), the spur-thighed tortoises displaying the highest rate (two subspecies added, n=7, 13%, Pr=30%). This species difference was observed among free-ranging tortoises (χ2= 15.76; df=3; p<0.01), but not among captive (χ2= 4.61; df=3; p=0.20) or vagrant tortoises (χ2= 2.44; df=1; p=0.11). There was no significant effect of the site on the prevalence of Mycoplasma in free ranging tortoises (χ2= 21.76; df=18; p=0.24. Table3). Focusing on WHT (wild, captive and vagrant pooled) tested positive for mycoplasma, there was no significant difference of the prevalence between the sexes (female’s Pr=9.7%, 23/238; male’s Pr=7.9%, 18/227; χ2= 0.25; df=1; p=0.62), in contrast to the herpes infection which occurred mainly in females. No individual was tested positive for both pathogens.

Considering the whole sample, 28 individuals showed clinical symptoms of URTD; 4 displayed palpebral oedema, 1 with sunken eyes, 8 strong nasal discharges, and 14 abnormal mucus color (black, white or yellow), 1 exhibited abnormal mucus color + nasal discharge. Among these 28 animals, 3 were tested positive for mycoplasma (2 with nasal discharges, one with palpebral oedema) and none for TeHV. The general condition of the tortoises was apparently normal otherwise and none exhibited ocular discharge.

## Discussion

This study reveals the first cases of herpesvirus infection and considerably extends knowledge of mycoplasma infection occurrence (only one case previously known; Mathes et al., 2001) in tortoises tested in natural populations in Europe (3 sub-populations for TeHV, 15 sub-populations for mycoplasma). These issues come sharply into focus when result obtained in captive and vagrant tortoises, native and exotic, are considered.

In this study, the high number of exotic tortoises observed vagrant (e.g., isolated individuals walking nearby private properties) or settled in WHT natural populations illustrate that uncontrolled introduction of exotic pet tortoises is an ongoing process. Chelonian pet trades are intensifying worldwide (Stanford et al., 2020), including European tortoises with massive exports of EHT and spur-thighed tortoises from Turkey, Balkan countries or North Africa to France, Spain and Italy (Ljubisavljević et al., 2011). Our results reveal substantial infection rates by mycoplasma both in captive and vagrant individuals of different tortoise species sampled in the distribution range of the native species (WHT). This situation poses the problem of possible transmission from captive toward free-ranging tortoises, and perhaps conversely when free ranging WHT are illegally collected and brought into captivity.

TeHV was not detected in captivity. The prevalence of mycoplasma infection was relatively similar in free-ranging and captive tortoises (Pr=8.7% and Pr=9.9% respectively). Among pet tortoises, however, infection rate was higher in exotic species and hybrids (22.2%, 4/18) compared to captive WTH (7.7%, 5/65). In addition, high levels of infection were observed in exotic spur-thighed tortoises, both in captivity (18.2%, 2/11) and among individuals introduced in the field (44%, 4/9) compared to those observed in free-ranging WHT populations (8.7%, 34/391). This disequilibrium between exotic and native tortoise’s infection levels suggests that a common exotic pet species may represent an important reservoir of mycoplasma. Wall-less mycoplasma were thought to be highly fragile; but the ability of some species to form resistant biofilms in vitro prompted a reconsideration of their ability to survive in the environment (McAuliffe et al., 2006). If M. agassizii would be able to form resistant biofilms in the environment, this may contribute to its persistent circulation even under low tortoise’s population densities. Although preliminary, these results reinforce the notion that pet tortoises, and thus international trade of chelonians, may represent a health threat to native tortoises.

Finding that free-ranging tortoises can be infected by TeHV and mycoplasma in Europe, although worrying, is not fully surprising. Negative SN tests for TeHV despite PCR positive tests suggest possible reactivation or recent contact with the pathogen that has not yet triggered immunity. Herpesvirus is a virus with a low environmental persistence capacity; transmission requires direct physical contact between tortoises (Marschang 2019). Numerous assessments performed in captive tortoises maintained in cages or in outdoor enclosures (inaccurately categorized as “wild” or “free-living” individuals by various authors, leading to misleading conclusions) showed that many individuals belonging to different lineages were infected by various pathogens involved in URTD. For example, herpesvirus was detected by PCR in 16.3% of individuals from different tortoise species maintained in captivity in Belgium; a country voided of native chelonian however (Martel et al., 2009). Kolesnik et al. (2017) tested by PCR 1,015 captive tortoises and terrapins originating from different continents and kept captive (e.g. in cage, enclosure, park, garden) in different European countries; 42.1% were infected by mycoplasma and 8.0% were infected by herpesviruses. In Turkey, among 272 spur-thighed tortoises caught in the wild and kept in captivity (sometimes during years) to supply a breeding program designed to reinforce native populations, seroprevalences (SN test) were as follow: 37% positive for herpesvirus serotype-I, 5.5% for serotype-II, and 5.2% for picorna-like ‘X’ virus and 4.9% for reovirus (Marschang and Schneider, 2007). Broadly similar PCR result of 36.7% was obtained with mycoplasma (Lecis et al., 2011).

Herpesvirus infections have been documented more than 40 years ago in American and in European chelonians (Origgi, 2012). However, previous assessments performed two decades ago in tortoises actually sampled in the field in France and Morocco (*Testudo hermanni, T. graeca*) failed to detect herpesvirus (Mathes et al., 2001; Mathes pers. com.). A situation that contrasts with well-documented infections in free-ranging chelonians in North America (Berish et al., 2000; Jacobson et al., 2012; Kane et al., 2017; Weitzman et al., 2017; Lindemann et al., 2019; Orton et al., 2020).

Further studies are needed to determine to what extent the presence of URTD pathogens results from recent transmissions from pet to wild tortoises in Europe, versus insufficient investigations in natural settings. Unfortunately, monitoring infection prevalence and severity is difficult in the field. Underestimations are likely because infected animals that may die are quickly removed prior to sampling. Further, weakened sick tortoises may well remain concealed and may not be easily detected in the field. This could explain the absence of co-infected individuals in our results.

### Infection risks and diseases

Lack of severe clinical signs should be treated with caution. Silent TeHV and Mycoplasma infections have been reported in tortoises (Orrigi, 2012; Withfield et al., 2018). Multiple strains of pathogens circulate worldwide (Salinas et al., 2011; Jacobson et al., 2014; Kolesnik et al., 2017), they continuously evolve while pathology and coinfections risks increase with pathogen diversity (Kari et al., 2008). Thus, constant introductions of infected pets into natural populations represent a threat through the emergence of infectious diseases, especially if exotic tortoises tolerate and thus carry pathogens that can cause outbreaks in less-tolerant native species (Jacobson et al., 2014; Whitfield et al., 2018; DiGeronimo, 2019; Goessling et al., 2019). Environmental factors, like seasonal variations of immunity and physiological stress can perturb equilibriums with pathogens, tipping the balance toward diseases; especially in case of multiple infections (Sandmeier et al., 2013; Goessling et al., 2016).

In European tortoises, the presence of herpesvirus and the high prevalence of mycoplasma in natural populations were only suspected prior to the current study. But the paucity of information regarding possible health impact of TeHV, in combination with mycoplasma, along with the spectrum of emerging serious diseases caused by picornavirus in captive Hermann’s tortoises in Spain for example (Martinez-Silvestre et al., 2020), prompts for further studies.

### Perspectives and recommendations

French continental populations of Hermann’s tortoise represent relicts of the distribution of a previously widespread and abundant species. It is crucial to not add infectious burden to the multitude of existing threats (e.g., habitat fragmentation, sprawling urbanization, frequent fires, illegal harvesting, wild-boars, dogs). Both the prevalence and demographic impact of URTD in the field should be carefully monitored. Pet owners should be informed that reproducing tortoises in captivity and releasing individuals in the field may threaten wild populations. Exotic tortoises should be removed from native populations. Long-term mark-recapture surveys should be coupled with health checking, including monitoring of most deleterious strains of TeHV (Gandar et al., 2015). Sanitary protocols are needed to handle free-ranging tortoises; nasal and oral secretions, feces, sperm and even urine can be contaminating (Origgi et al., 2012). Similarly, strict protocols are needed during conservation translocations; notably because individuals often travel long distances before settling (Pille et al., 2018), while stress associated with release may promote virus reactivation from latent infection (Griffith, 1993; Martel et al., 2009; Jacobson et al., 2012). Furthermore, captive individuals carrying a wide range of pathogens and parasites represent additional risks to URDT transmission and should be treated specifically (Ahne 1993; Chávarri et al., 2012).

## Data accessibility

Data are available online: Doi: 10.5281/zenodo.4467755

## Acknowledgements

We thank Silvia Blahak (Detmold) for herpesvirus and mycoplasm testing. We thank Concha Agero, from the Reserve National de la Plaine des Maures; Pierre Lacosse, Camille Casteran and Bryan Teissier from the Port-Cros National Park; and Antoine Catard from the Conservatoire des Espaces Naturels de PACA, for providing us authorization to access their property and help in the field. This study was funded by “La Fondation Klaus Zegarski pour la conservation et protection des tortues méditerranéennes”, and the DREAL-PACA. This project was conducted under the permits delivered by prefectoral authorities on July, 13th 2012 and February, 26th 2013. Version 4 of this preprint has been peer-reviewed and recommended by Peer Community In Zoology (https://doi.org/10.24072/pci.zool.100007).

